# Measurement of Oxidative Stress in Huh-7 Cell Line to Determine the Effectiveness of PNPLA3 Targeted Gene Therapy for Mitigation of Lipid and Alcohol Induced Oxidative Stress in the Liver

**DOI:** 10.1101/2023.05.20.539653

**Authors:** Jihoon Kwon, Elena Ivanovna Budyak

## Abstract

NAFLD is a condition of increased buildup of fat in the liver, causing lipotoxicity that can manifest to cirrhosis, steatosis and fibrosis, which can cause significant and eventually irreversible damage to the liver. A key gene associated is the PNPLA3 I148M variant, which has been shown to display lipogenesis functionality. However, the role of PNPLA3 in increased ROS formation is debated. Moreover, there are no studies determining correlation between the variant and alcohol-induced oxidative stress. This project determines the following mechanistic functions of the PNPLA I148M variant to test for the efficiency of PNPLA3 gene therapy for patients with fatty liver disease and alcohol liver disease.

There is a strong correlation between lipid induced oxidative stress and PNPLA3 148M variant. DCFDA shows increased concentration of H_2_O_2_ for PNPLA3 148M overexpressed cell lines for both ethanol and FFA treatment groups. Moreover, there is a statistical correlation between PNPLA3 148M overexpressed cell lines and increased mitochondrial oxidative stress by MitoSOX cellular ROS analysis methods. This study confirmed the significant decrease in oxidative stress levels for 148I variant overexpressed cell lines and proved the efficiency of PNPLA3 targeted gene therapy to NAFLD patients and suggested the potential use of the therapeutic method to patients with ALD. To ensure gene therapeutic effectiveness for patients with NAFLD and alcoholic liver diseases, further experiments may be needed to verify molecular pathways of ROS formation by PNPLA3 with qPCR analysis.

## Introduction

Non-alcoholic fatty liver disease (NAFLD) is a disorder in which large amounts of fat build up in the liver. NAFLD can manifest into non-alcoholic steatohepatitis in which the liver becomes inflamed and damaged.^25^ NAFLD is a big contributor to morbidity and mortality worldwide because of the high rate of heart, liver, and cancerous complications associated with it. NAFLD is also becoming the leading cause of end stage liver disease and liver transplants.^4^ A key gene associated with NAFLD is patatin-like phospholipase domain containing protein 3 gene (PNPLA3). It is a gene located on chromosome 22, and is associated with lipid metabolism.^1^ Moreover, NAFLD is can induce lipotoxicity in the liver, in which too much lipid accumulation can cause a toxic environment in the liver and leads to oxidative stress that results in inflammation of liver tissues and damage in liver cells.^25^

When lipotoxicity is induced in the liver, the PNPLA3 gene is mutated from isoleucine to methionine attached on position 148 of the gene strand, which is a single nucleotide polymorphism (SNP). This mutation gives rise to the 148I and 148M variants of the PNPLA3 gene, and can be acquired through environmental factors like lipotoxicity, or can be congenitally acquired.^5^ This mutation has been studied, and it has been determined that this mutation increases the retention of lipids in the hepatocytes, leading to exacerbation and positive feedback loop of lipid accumulation causing more damage through more lipid accumulation.^12^

Another profound effect that lipotoxicity can have on the liver is the mitochondrial depolarization process that initiates in the hepatocytes and is induced by reactive oxygen species (ROS).^29^ Mitochondrial depolarization leads to pores on the mitochondrial membrane, causing a fluid called Cytochrome C to leak out of the mitochondria and into the membrane. The Cytochrome C will then activate a cascade of events that will lead to apoptosis, a programmed cell death. Normally, this occurs naturally if the space within the cell is too crowded, or if a cell is infected by a pathogen. However, because lipotoxicity directly induces apoptosis, cell death starts at a faster rate than the rate at which the liver can replace itself, causing further damage to the liver. Moreover, mitochondria is a vital part of lipid metabolism as they synthesize ATP that is used for metabolism. Therefore, determining the condition of the mitochondria is important when studying the effects of oxidative stress on the liver cells.

Going back to the PNPLA3 gene, it codes for adiponutrin, a protein made with transcription and translation of PNPLA3 gene. Adiponutrins are found in fat cells and liver cells and function in lipid metabolism and reallocation of excess lipids into the adipocytes all throughout the body.^1^ However, when lipotoxicity in the liver is induced, 148M mutation occurs and the mutation impairs the PNPLA3 protein turnover process.^8^ This occurs due to the mutation impairing the functionalities of ubiquitin and autophagosomes. Ubiquitin is a regulatory protein that functions in apoptosis, protein processing, and DNA repair. Additionally, autophagy mediated degradation is the process in which autophagosomes take in proteins and organelles and then fuse with the lysosomes for degradation of the internal contents. Protein turnover is the replacement of older proteins that are dysfunctional and inactivated with newer and functioning ones by breaking down older proteins within the cells. However, the 148M mutation impairing the ubiquitin function and autophagy mediated degradation process leads to old and dysfunctional PNPLA3 proteins to accumulate in the liver, which further exacerbates the lipotoxicity condition.

NAFLD can affect anyone, regardless of age and genetic makeup. As Richard Lehner and Ariel D Quiroga, adjunct professors at University of Alberta Department of Cell Biology, states in their study that NAFLD can affect children as well as adults.^11^ Moreover, the genotypic difference can change the likelihood of a person acquiring the disease, however, any type of genetic makeup can cause NAFLD to develop. Nader Salari, an Associate Professor of Biostatistics at Kermanshah University of Medical Sciences, lists the percentage likelihood of someone acquiring the disease based on their genes. The professor concluded that “homozygous recessive genotype patients for PNPLA3 where the expression was minimal, their risk of developing NAFLD had 52% lower chance, heterozygous genotype had increased to 19% higher risk, and for homozygous dominant genotype, the percentage rose to 105%.” ^19^ This shows that the mutation is strongly associated with susceptibility to the development of the NAFLD, and it can affect anyone, regardless of their genotypic predisposition.

Even though NAFLD can be prevented by merely changing diet and eating less high fat foods, it nonetheless remains a highly prominent and problematic disease. This is due to the fact that lower income Americans eat unhealthier than the higher income Americans because of their lack of access to healthier foods.^2^ The lack of healthy food options for lower income Americans, along with the lack of healthcare access provided to them, makes NAFLD a problematic disease and highlights the urgency to develop a therapeutic method for the mitigation of damage caused by the disorder.

Basic Physiology of Fatty Liver Disease

Fats and cholesterol make up one of the main macromolecules found in the body. Cholesterol functions in maintaining the structure and fluidity of cell membranes. However, too much accumulation of cholesterol can cause many problems, including the development of NAFLD. Studies show that increased consumption of foods high in fat and cholesterol was linked with activating opioid and dopamine receptors in the brain.^4^ The changes in the reward center such that only the foods high in fat stimulate them will cause more accumulation of lipid in the liver due to the person consuming more high-fat foods which further exacerbates the conditions of NAFLD and leads to further damage in the liver, which is responsible for detoxifying and redirecting lipids into the adipocytes. Insulin is also a major contributor to lipid metabolism. It promotes adipose tissue to store lipid droplets and the hepatocytes to activate key regulators of lipogenesis that balances the level of fat accumulation in the liver.^6^ However, for patients with NAFLD, the resistance to insulin develops which increases the concentration of lipids flowing through the circulatory system, which will result in net increase in lipid accumulation in the liver.^6^

Pathway of Normal Alcohol Metabolism

Alcohol is metabolized by various enzymes like aldehyde dehydrogenase (ALDH), alcohol dehydrogenase (ADH), cytochrome P450 (CYP2E1). Ethanol is converted into acetaldehyde by ADH in the cytosol. Then, it is covered into acetic acid with ALDH_2_ in the mitochondria. And it is further broken down into acetyl-CoA chemical.^26^ Acetyl-CoA, then enters the citric acid cycle, which produces citric acid, yielding a relatively non-toxic chemical from ethanol.^28^ However, in some cases ALDH_2_ and CYP2E1 malfunction due to the drastic pH change from alcohol accumulation, acetaldehyde may not be converted into citric acid from the citric acid cycle. Acetaldehyde can lead to overproduction of ROS and impairs mitochondrial function. This not only damages the ATP production, it also causes oxidative stress in the liver that can cause apoptosis and toxicity. The pathway for alcohol metabolism is outlined below.

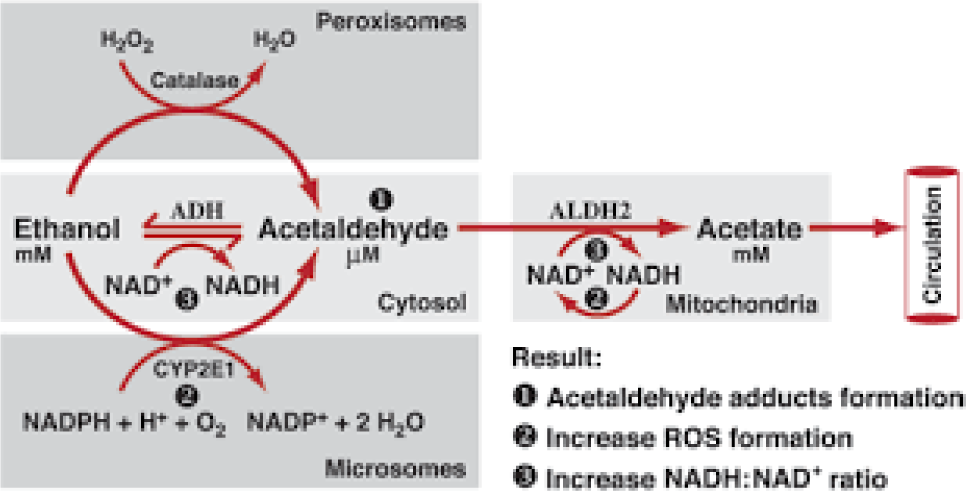

Image source: Szabo, G., & Saha, B. (2010). Alcohol’s Effect on Host Defense. Alcohol Research & Health, 33(1-2), 47-63. PMID: 23580036.

Role of Mitochondria in Lipid Metabolism

Mitochondria plays an integral part of lipid metabolism. ATP, which is produced in the mitochondrial matrix of the liver, is instrumental in the metabolism of lipids. Triglycerides that enter the liver are oxidized by the β-oxidation process. This process to turn fatty acids and triglycerides into acetyl CoA requires ATP.^13^ Therefore, any damage in the mitochondria will impair the production of ATP and therefore the metabolism of lipids is also impaired, so the maintenance of mitochondrial integrity is imperative for patients with NAFLD. Moreover, the structural integrity of mitochondrial membrane for the prevention of apoptosis is vital for the mitigation of oxidative damage. Therefore, to better understand how the PNPLA3 gene and the oxidative stress affects mitochondrias, MitoSOX analysis of mitochondrial oxidative damage was performed which will be outlined in the methodology section.

PNPLA3 Targeted Gene Therapy

There are several methods of gene therapy, one of them being gene inactivation. Inactivation of a disease-causing gene that is dysfunctional can be done with small interfering RNA (siRNA), short hairpin RNA (shRNA), and antisense oligonucleotides.^7^ Small interfering RNA are artificially synthesized and are used for silencing genes of interest. SiRNA binds with the gene of interest, and transfers it to a protein called AGO2, which then develops a cleavage in the passenger strand of the DNA.^16^ The target mRNA is cut and the gene is silenced. Another way that a gene can be silenced is by using shRNA, which uses a double stand of siRNA in order to clip the target gene, PNPLA3 in this case, and then silence the gene.^20^ Finally, antisense oligonucleotides can be used to silence the PNPLA3 gene by cleaving the RNA-DNA complex with the binding of itself with the RNA. Antisense oligonucleotide is a single-stranded deoxyribonucleotide, and the form of antisense oligonucleotide can allow it to bind to the RNA so that the RNA that was transcribed into RNA is prevented from translating into proteins because of the antisense oligonucleotide binding to it.^7^ These are the methods in which genes can be silenced. Another method of gene therapy is the injection of healthy genes into the body, like the 148I variant. This is done in two ways: in vivo, and ex vivo. In vivo method includes the injection of gene filled fluid into the body intravenously. In contrast, ex vivo method includes the extraction of tissue cells from the body and then the gene editing occurs in the lab before the cell is inserted back into the body for the purpose of multiplying throughout the body and spreading the healthy gene to other cells.

### Literature Review of Relevant Sources

It is widely known throughout the scientific community that the PNPLA3 gene is associated with NAFLD and NASH. However, an exact mechanism of how the PNPLA3 148M variant is associated with increased risk of chronic liver disease development is still debated.^15^ Moreover, Dr. Dong states that many genotype wide association studies (GWAS) of different racial groups have shown that the 148M mutation of PNPLA3 is associated with lipid accumulation, NAFLD, and NASH regardless of alcohol consumption.^8^ Dong also states that because the PNPLA3 148M variant is prevalent among most populations, it is necessary to develop a therapeutic method that targets this genetic mutation.^8^ This shows that there is a need for a PNPLA3 targeted gene therapy for patients with NASH and NAFLD. Furthermore, Dong states in his abstract that “with additional mechanistic studies, novel therapeutic strategies are expected to be developed for the treatment of the PNPLA3 148M variant-associated chronic liver diseases in the near future.”^8^ This is a clear gap and lack of knowledge in this field regarding the functionality of PNPLA3 that, if further investigated, would provide the additional value in the development of PNPLA3 gene therapy as a useful therapeutic target.

Another article by Francesca V. Bruschi and various other professors in Austria investigated the mechanism of PNPLA3. The method they used in Bruschi’s paper had many similar aspects for determining the mechanism of PNPLA3 with the research method used in this paper. In this article the authors use Huh-7 cell line, immunofluorescent imaging technique, and palmitic acid to induce lipotoxicity. Moreover, they also use liver samples of patients to study PNPLA3 gene’s role in liver fibrosis. Through this research, they are able to conclude that PNPLA3 increases during lipotoxicity in the liver caused by the liver cells obtained in the clinic, as well as the cell lines.^3^ However, with the immunofluorescence imaging, the researchers only determine the concentration of PNPLA3 to reach their conclusion.

The research was designed to build upon what Francesca and other professors in Austria conducted in their study. They have established that PNPLA3 gene expression increases in the liver with the addition of palmitic acid or the occurrence of non-alcoholic liver disease. To add to the current body of scientific knowledge surrounding the functionality of the PNPLA3 gene, Cellular ROS analysis was performed to quantify the amount of oxidative stress that has occurred due to the overexpression of PNPLA3 variants. Dr. Solomon highlights the significance of ROS by pointing out that ROS is the sole causative agent for inflammation and cell death in the organ.^22^ Therefore, analyzing the effect that PNPLA3 has on the formation of ROS will better support, or discourage the notion that PNPLA3 is an excellent therapeutic target for patients with lipotoxicity from NASH and NAFLD.

Several articles above indicate the association between PNPLA3 and liver diseases like NAFLD and NASH. Theoretically, PNPLA3 targeted gene therapy for patients with liver disease will be a beneficial therapeutic treatment method. To investigate this gap, the following question was formed: “Using PNPLA3 overexpression, to what extent would targeted PNPLA3 gene therapy mitigates oxidative stress levels induced by lipids and alcohol.” The hypothesis is that the Wild Type group with no transfection will display the lowest oxidative stress levels while the 148M overexpressed cell line will exhibit the most. Overall, studies show that PNPLA3 expression increases in the liver and cell lines when lipotoxicity is induced in those cell lines or in patient’s livers by palmitic acid or NAFLD. However, little is known about the ROS activity in patients with PNPLA3 overexpression. Determining this information is the purpose of this study

## Methods

The method used in this study was a gain-of-function experimental research approach. In order to establish correlation between the PNPLA3 gene variants and their oxidative stress levels, a gain-of-function approach was used to amplify the effects of the 148I and 148M variants to determine a clearer correlation between the PNPLA3 gene variants and oxidative stress. To determine the efficiency of the PNPLA3 targeted gene therapy, Huh-7 cell lines were used. Huh-7 cells, which are immortalized liver cells, were chosen because they were the best alternative to human livers and much more ethical to conduct manipulations on than human livers or primary hepatocytes harvested from mice. Moreover, Huh-7 cell lines are also frequently used as a precursor by clinical scientists to collect in-vitro data before they proceed with clinical trials, especially with the studies in lipoprotein metabolisms.^14^ The Huh-7 cell lines were cultured in an incubator where they were kept in 37 degrees celsius and exposed to constant flow of carbon dioxide and oxygen to simulate what liver cells experience in a live body. Other equipments that were used for this experiment were the following: palmitic acid, ethanol, anti-PNPLA3 antibodies, DAPI staining, BODIPY staining, DCFDA dyes, MitoSOX superoxide indicator, and fluorescence microscopy.

Huh 7 Cell Line Establishment, Transfection Done by Plasmid and E. coli

In order to observe the effects of both lipid and alcohol on liver cells, nine Huh-7 cell lines were used. For Huh-7 cell line establishment, The Huh-7 cells were placed in nine 35 mm round glass bottom dishes. Then, they were cultured in an incubator with the conditions mentioned above until the cell density was 5 × 10^4^ cells/mL. Then the nine cell lines were divided into three groups: normal, PNPLA3 148I overexpressed, and PNPLA3 148M overexpressed. This was done with transfection. Transfection is a way to introduce foreign nucleic acids into eukaryotic cells.^23^ This was done by combining the plasmid containing PNPLA3 148M variant or 148I variant with an aqueous solution with E. coli. Then, the solution was incubated in ice and warm water alternatively. During this time, the E. coli bacteria takes in the plasmid from the solution and integrates it into themselves in a process called bacterial transformation. Therefore, the PNPLA3 genes multiplied along with the E. coli bacteria. This PNPLA3 gene was then extracted with a centrifuge that uses high velocity movement to separate plasmid and E. coli, yielding the fluid of PNPLA3 gene.^9^ However, direct application of this fluid to the cell line would not work, as cells generally do not absorb DNA clusters. This problem can be solved by adding lipid coated vesicles around the DNA, called Lipofectamine. When PNPLA3 gene filled fluid and Lipofectamine were both applied to the cell line, it was left for two to three days for most of the DNA to diffuse into cultured Huh-7 cell line. The pathway for gene introduction into the cell by Lipofectamine is shown in the diagram below

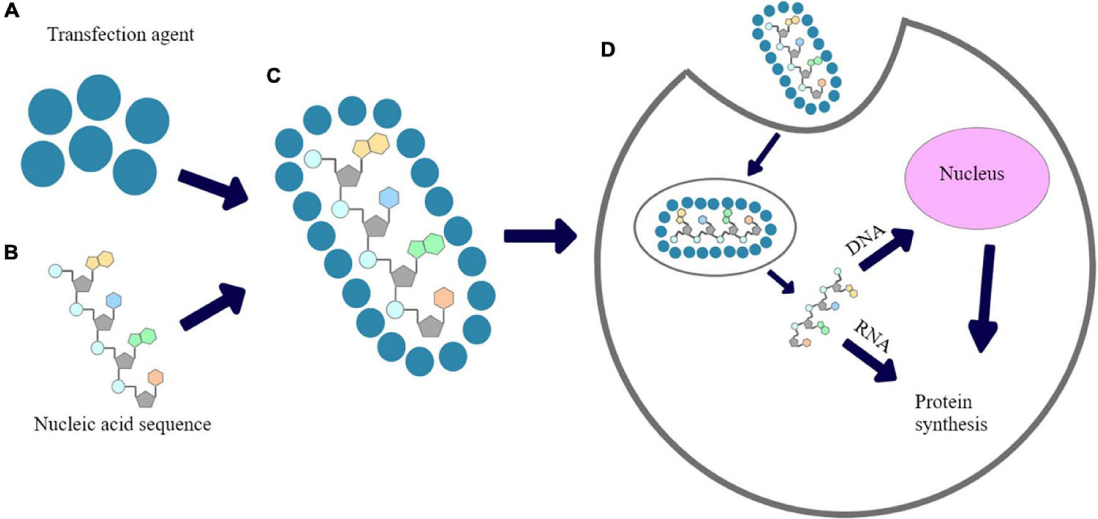

Images source: Jafarizadeh-Malmiri H, Keyhani A, Mohammad-Abadi M, et al. Microbial production of polyhydroxyalkanoates (PHAs) and its copolymers using food waste streams: a review. Front Bioeng Biotechnol. 2021;9:701031. doi: 10.3389/fbioe.2021.701031.

Genotyping done with DAPI, Bodipy, antiPNPLA3 antibodies

To confirm that the transfection was successful, a series of stainings were done to identify the overexpression of PNPLA3 genes, presence of liver cells, and accumulation of lipid droplets. All three types of the transfected cell lines were applied with 2.5% normal horse serum to clean any debris and sterilize it before applying antibodies and stains. Then, the primary antibodies consisting of goat anti-PNPLA antibodies, which can detect both the 148M and 148I variant proteins, were applied to the cell lines. Then, the cell lines were incubated at 4 °C overnight. Then, it was washed with phosphate-buffered saline solution with a low-concentration detergent solution twice and sterilized phosphate-buffered saline once. These solutions are non-toxic and contain the optimal pH value and concentration of ions that cleanses the excess anti-PNPLA3 antibodies that did not bind to the proteins without any cellular damage or shrinkage of the cells. Then, the cell lines were applied with secondary antibodies to detect the site of binding and emit fluorescence with microscopy imaging. The secondary antibodies were attached with Alexa 647 fluorescence that can be excited when met certain lightwaves and display fluorescence. Then, the cell lines were applied with Bodipy staining of 1 µM for detection of lipid droplet area and it was washed again with the solutions to get rid of all the excess Bodipy staining.^17^ It was then applied with DAPI staining for nucleus detection, in order to confirm the presence of liver cells and quantify the amount of them in each image.^24^ The cell lines were washed systematically with the solutions and then mounted to the ZEISS microscopes that emitted the specific frequency of waves to create a fluorescence image of each type of cell line.

The first three sets of images below show the representative cell line for each transfection group. Anti-PNPLA3 antibodies were applied for detection of proteins made by the PNPLA3 gene. The overexpressed cell lines exhibited the most redness which confirms transfection. Then, Bodipy staining was used to detect lipid droplets. This was to differentiate between the 148I and 148M overexpressed cell lines. Because the 148M variant functions in lipogenesis, this could be confirmed by comparing the presence of lipid droplets between the overexpressed cell lines. The green fluorescence was most prominent in the PNPLA3 148M overexpressed cell line, confirming the 148M gene presence in that cell line. Finally, DAPI staining, which detects nuclei, showed blue clumps in all of the cell lines, which confirms that the liver cells are present. Refer to Fig. 1.

**Figure 1.**
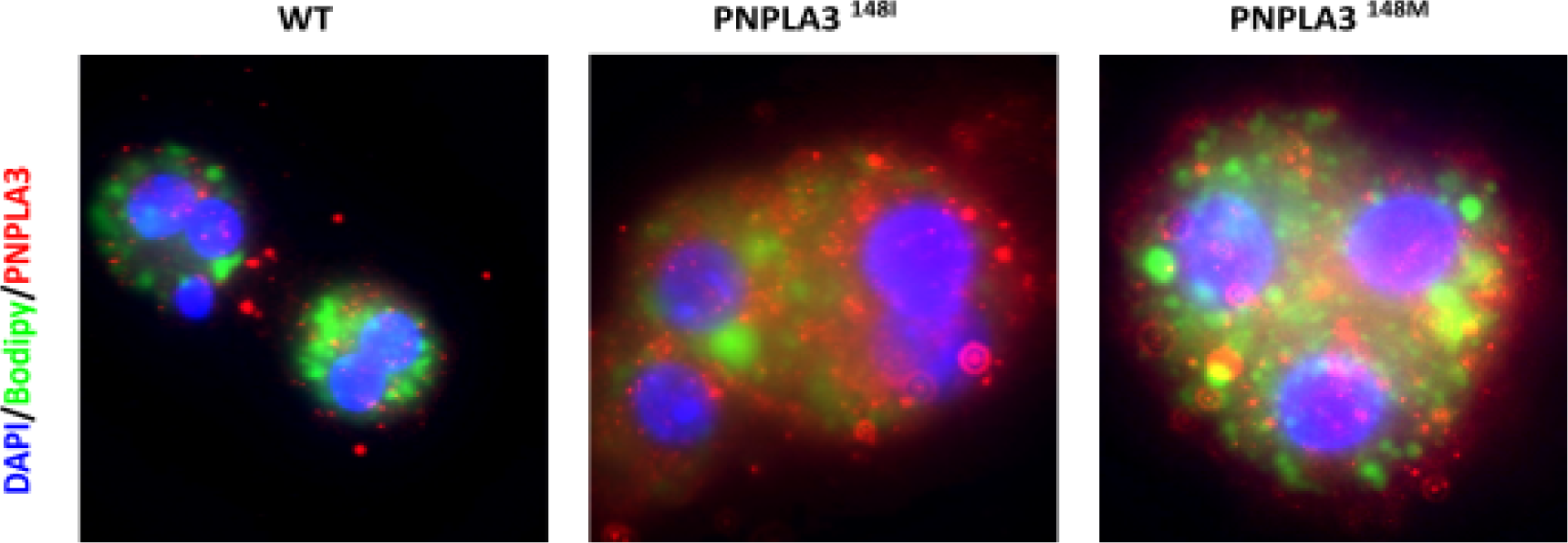
Genotyping fluorescence imaging data: the PNPLA3 overexpressed cell line had increased red expression and PNPLA3 148M overexpressed cell line showed increased green expression, confirming gene transfection

Lipotoxicity in the Liver Induced by Acids, Alcoholic Liver Established by Ethanol

After all the cell lines were divided into groups according to the gene variant transfection and confirmed with genotyping, they were further divided into groups according to the treatment that they will receive with random assignment method in order to investigate the effect that the lipid and alcohol has on the liver. One of those groups was not given any treatment to establish a baseline for the oxidative stress while the other two groups were given FFA and Ethanol treatment. For FFA treatment, both palmitic acid and oleic acid were used. Palmitic acid is the primary source that causes lipotoxicity, however, this can also cause cells to burst.^18^ Oleic acid is a naturally occurring and non-toxic form of fatty acid, which does not induce lipotoxicity. In order to minimize cell death while still inducing lipotoxicity in the cell lines, palmitic and oleic acid were combined in a 1:3 ratio to create a diluted palmitic acid solution of 320 µM. The Ethanol group was treated with 10 mM of ethanol, a compound commonly found in alcoholic beverages.^28^ These cell lines were then incubated for 48 hrs for the lipids and alcohol to take effect before being analyzed for oxidative stress.

MitoSOX cellular ROS analysis

The first cellular ROS analysis using MitoSOX indicators that detect mitochondrial oxidative stress. Detection of mitochondrial oxidative stress is important because lipid metabolism is done mainly with ATP produced in mitochondria. Oxidative stress in the mitochondria will exacerbate the lipid accumulation because mitochondria cannot make the ATP for lipid metabolism, which will exacerbate the fibrosis and cirrhosis diseases.^27^ 5 µM of MitoSOX staining was applied to the cell line which marks the mitochondrial oxidative stress. After around 10 minutes, the cell lines were washed with 10mM of PBS solution of 7.3 pH level. The cell lines were fixed and prevented from decay and dying off with 4% concentrated paraformaldehyde (PFA). With the MitoSOX applied and cell fixed wavelengths were projected onto the cell with ZEISS microscopy. MitoSOX absorbs at 396 nm wavelength lights and emits fluorescence of 610 nm in response to the absorption. Red light emission from the MitoSOX detects mitochondrial oxidative stress.^10^ When combined with the baseline blue light emission from normal mitochondria, oxidative stress shows up as purple oval like shapes in the fluorescence images.

The results for MitoSOX staining for the 9 Huh-7 cell lines are shown below. MitoSOX marker detects mitochondrial oxidative damage and emits red light. Based on preliminary observation, the PNPLA3 148M cell line displays not only increased redness but also has reduced numbers of mitochondria present compared with the WT and 148I cell lines in the same treatment groups. Refer to Fig. 2.

**Figure 2.**
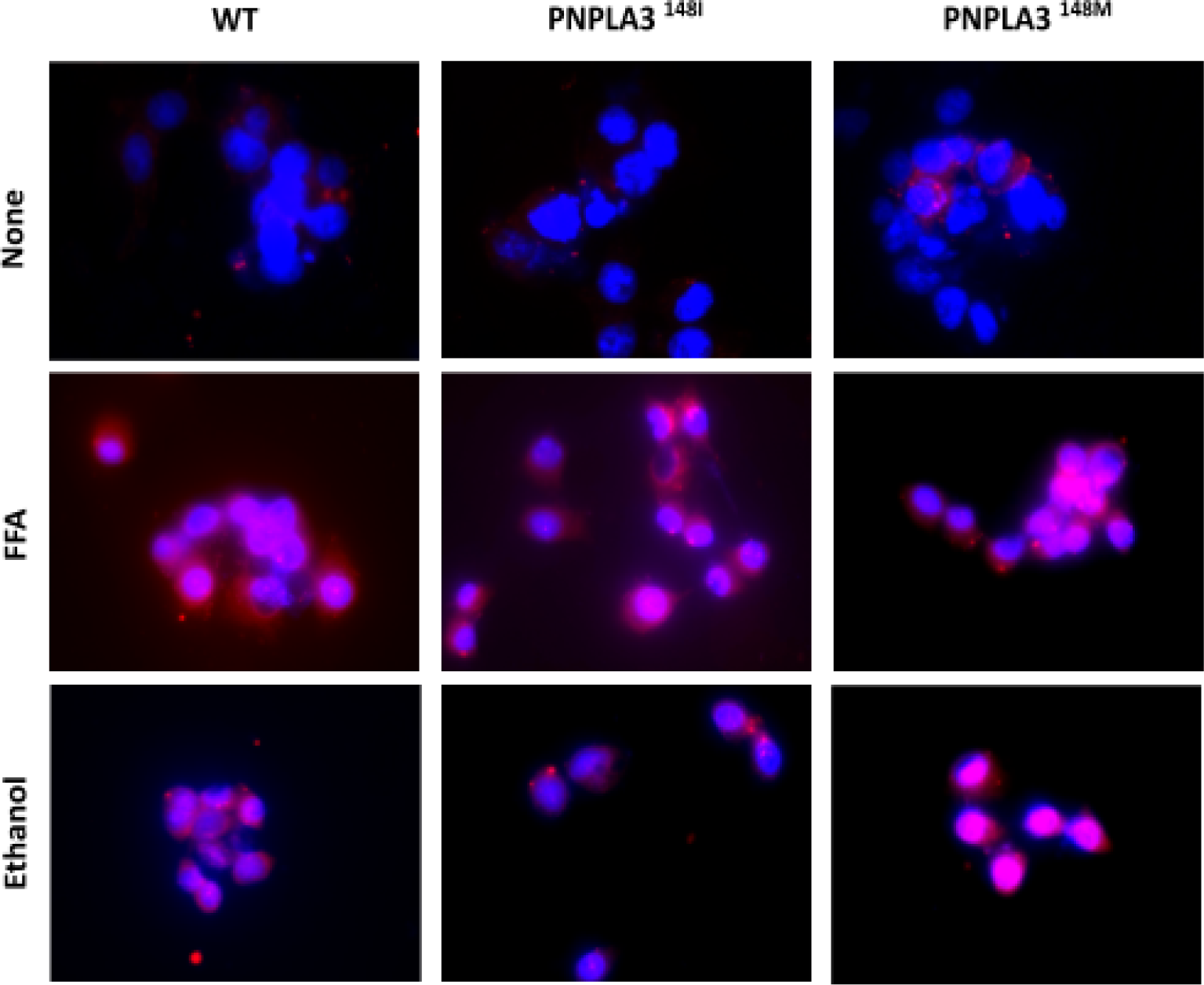
MitoSOX fluorescence imaging data: 148M overexpressed cell line displayed more red fluorescence and less mitochondria compared to other groups

DCFDA cellular ROS analysis

In addition, cellular ROS assay using the DCFDA staining method was performed. This was used to determine the concentration of ROS for the cell lines. Dichlorofluorescein diacetate (DCFDA) was applied to both of the cell lines, and when DCFDA makes contact with H2O2, then an oxidation reaction occurs between these two chemicals that yields the product dichlorofluorescein, which is highly fluorescent and produces greenish yellow lights with the absorption of the wavelengths of 498 nanometers and 522 nanometers. With DCFDA staining on both cell lines, determination of both the presence and the concentration of H2O2, a prominent indicator that damage in the liver has been done, is possible.

The results for DCFDA staining for the 9 Huh-7 cell lines are shown below. DCFDA detects H_2_O_2_ in the cell lines and emits yellowish green fluorescence when they do so. Based on preliminary observation, there is the most amount of H_2_O_2_ in 148M cell lines. Moreover, there is a decrease in the hydrogen peroxide formation for the 148I cell line as compared to the wild type, which further strengthens the claims that support the usage of PNPLA3 gene therapy. Refer to Fig. 3.

**Figure 3.**
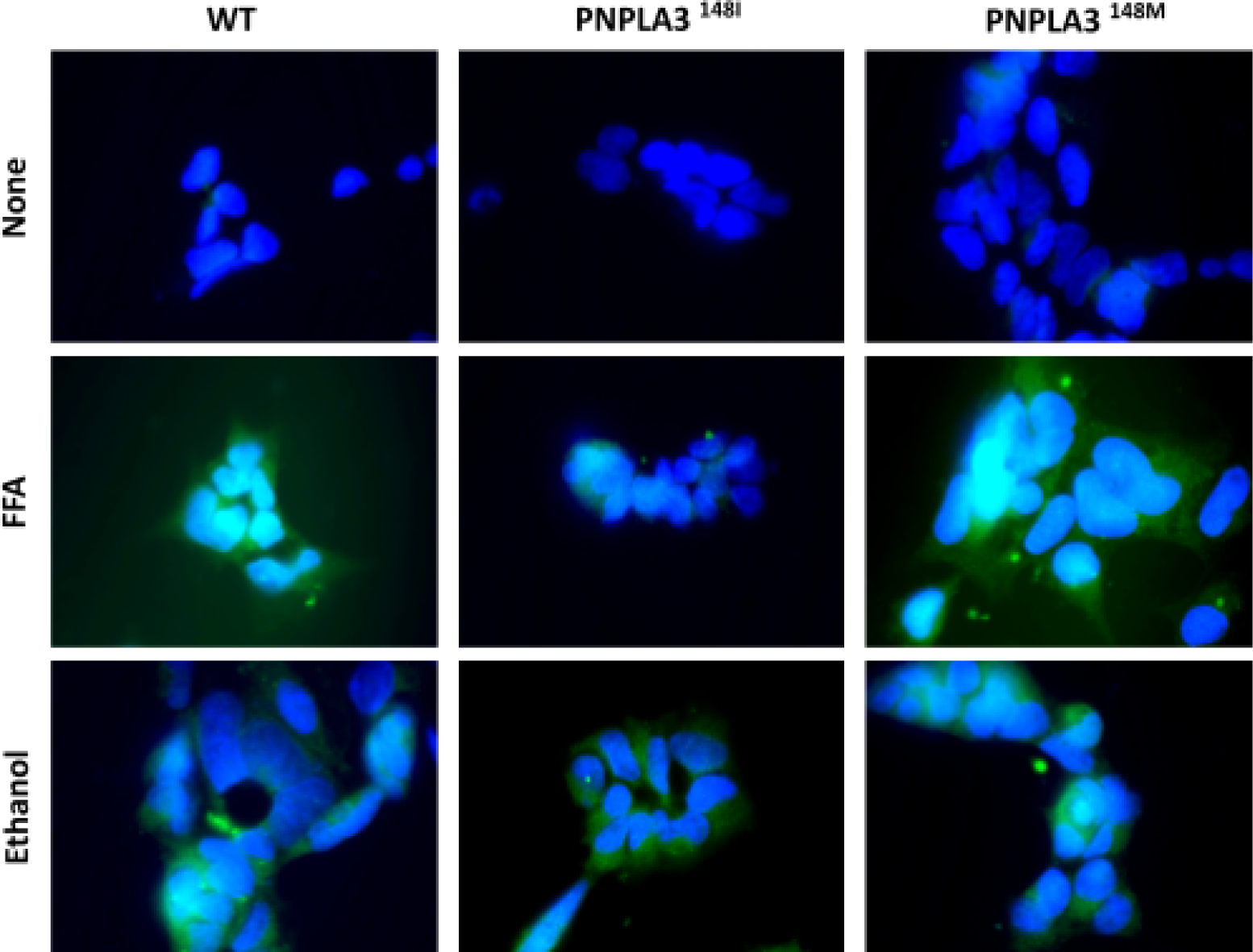
DCFDA fluorescence imaging data: 148M group displays the most among of greenish yellow fluorescence as compared to other groups

### Ethics

As cancerous immortalized cell lines derived from a human being were used, ethics needs to be considered. The origins of the Huh-7 cell line is well known in the scientific community, and the person donated his liver cell and consented to its use in clinical research. Nonetheless, to maintain ethicality and confidentiality, no personal identifying information was released about the person whom the Huh-7 cell line was derived from in order to maintain ethicality and anonymity of all parties involved.

## Results

Each of these 9 images were analyzed with two trials of mean fluorescence intensity and were organized into the data tables below. Refer to Fig. 4. Fluorescence intensity is measured by detecting photon emission of the ROS stains in the cell lines. When laser beam is passed through the Huh-7 cell lines, the photons are emitted and are detected by the photomultiplier tube (PMT) and then registered in real time histogram with the peaks representing the number of photon emission.^21^ The peaks are then averaged to yield a fluorescence intensity data for one of the cell lines. Two trials were done for determining the fluorescence intensity for both MitoSOX and DCFDA cellular ROS analysis images, and the average of the two trials were calculated, as well as the averages of the 4 total DCFDA and MitoSOX mean fluorescence intensity values. These 4 data points for each of the 9 cell lines, along with the mean fluorescence and statistical significance lines were then put into a bar graph format. The data tables showing the fluorescence intensities for the ROS analysis and the bar graph are shown below. Refer to Fig. 5.

**Figure 4.**
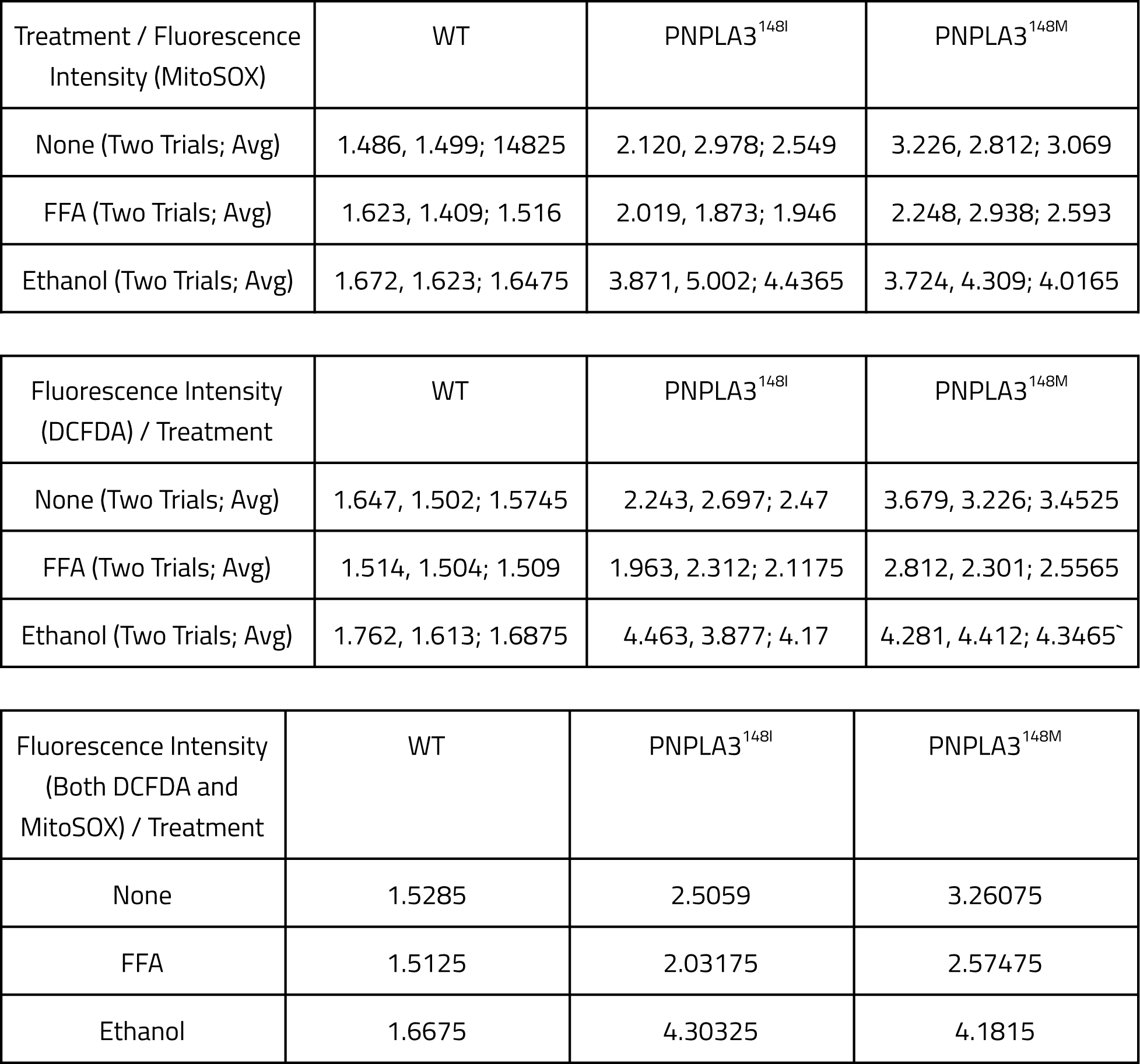
Fluorescence Intensity Unit Data Table

**Figure 5.**
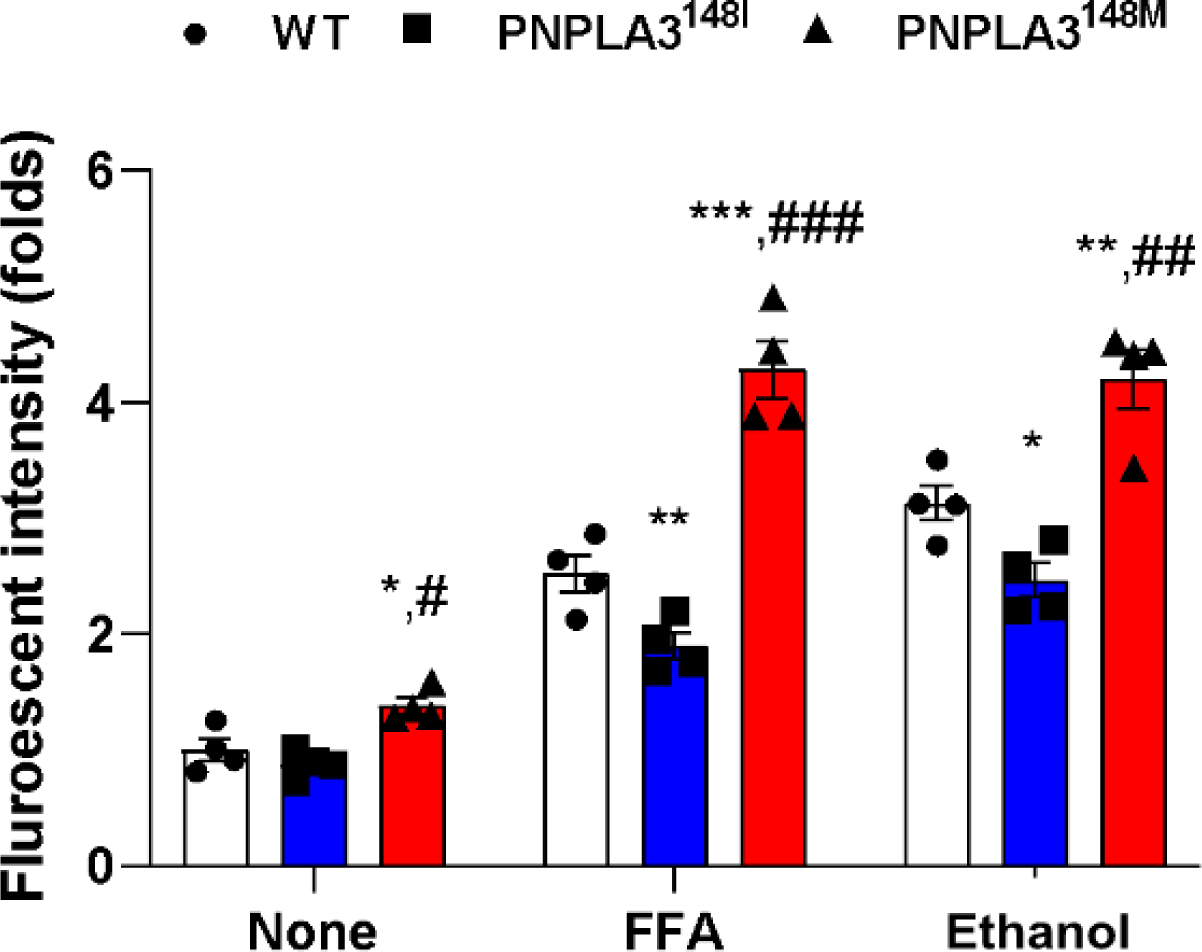
The fluorescence intensity as shown in this bar graphs is organized into folds, which is in a logarithmic scale, therefore, fluorescent intensity increase from 0 to 2 would be 10^2^ percent increase, and from 0 to 4 would be 10^4^ percent increase. P values were also added, * on 148I variant bars show the p value between WT and 148I variant, * on 148M variant bars show p value between WT and 148M variant, and # shows the p value between 148I and 148 M variant. * or # represents p value less than 0.05, ** or ## represents p value less than 0.01, and *** or ### represents p value less than 0.001

## Discussion

Each set of nine images shows each of the three cell lines with different gene transfection that were treated in three different ways. MitoSOX marker detects mitochondrial oxidative damage and emits red light. Based on preliminary observation, the PNPLA3 148M cell line displays not only increased redness but also has reduced numbers of mitochondria present compared with the other cell lines. DCFDA detects H_2_O_2_ in the cell lines and emits yellowish green fluorescence when they do so. Based on preliminary observation, there seems to be the most amount of H_2_O_2_ in 148M cell lines. Moreover, there seems to be a decrease in the hydrogen peroxide formation for the 148I cell line as compared to the wild type, which further strengthens the claims that support the usage of PNPLA3 gene therapy.

Upon statistical analysis of mean fluorescence values, it was clear that there were most oxidative stress levels and damage in the PNPLA3 148M cell lines across the board. Moreover, the cell lines with PNPLA3 148I cell lines had the least fluorescence, oxidative stress, and damage. This corresponded with the initial hypothesis of this research regarding the functionality of PNPLA3 gene, which was that the lipogenesis function of the 148M variant will correlate with more oxidative stress as there will be increased lipid accumulation in the liver, and the lipid metabolism function of the 148I variant will correlate with less oxidative stress.

The gain-of-function research was done through gene overexpression to determine the effects of PNPLA3 gene clearly. As compared to the data that gene knockout (KO) method produced, the gain of the function research model produced clear differences in oxidative stress levels between the cell line groups. The differences in the fluorescent intensity levels were very clear. The difference between WT and PNPLA3 148M overexpressed cell lines that were treated with FFA was 1.8 ± 0.03, and for the Ethanol treatment group, the difference was 0.92 ± 0.03. These statistics show that the PNPLA3 targeted gene therapy would decrease oxidative stress by 42% for patients with NAFLD and decrease oxidative stress by around 22.1% for patients with ALD. This shows the benefit of PNPLA3 targeted gene therapy that inactivates the PNPLA3 148M gene for patients with NAFLD and ALD. However, additional benefit was shown for 148I overexpression. The difference between WT and PNPLA3 148M overexpressed cell lines that were treated with FFA was 2.27 ± 0.03, and for the Ethanol treatment group, the difference was 1.61 ± 0.03. These statistics show that PNPLA3 targeted gene therapy which includes not only the inactivation of PNPLA3 148M variant but also the injection of the healthy 148I variant will decrease oxidative stress by 53% in NAFLD patients and will decrease oxidative stress by 39% in ALD patients. Furthermore, because of the oxidative stress Other previous research indicated that the PNPLA3 148M variant was highly prominent among patients with NAFLD.^19^ Moreover, other researchers determined that the PNPLA3 gene increases with NAFLD and ALD.^8^ Through the developed methods, oxidative stress increase of the mutated variant of the PNPLA3 gene was determined. The result of this study shows that PNPLA3 targeted gene therapy that includes PNPLA3 148M inactivation and injection of 148I variant will be a very competitive and effective treatment method for patients with not only NAFLD but also ALD.

## Conclusion

The purpose of this study was to determine the effectiveness of PNPLA3 targeted gene therapy. Gain-of-function research was chosen to determine the oxidative stress level differences between the mutated variant of PNPLA3 and the physiological variant of PNPLA3. Gene transfection was performed on selected cell lines intended for PNPLA3 gene overexpression, followed by a series of genotyping staining to confirm that gene transfection was successful. After genotyping, MitoSOX and DCFDA cellular ROS were done in order to determine any potential differences between the oxidative stress levels in normal and mutated variants of PNPLA3 genes. Through immunofluorescence imaging analysis, it was determined that the PNPLA3 targeted gene therapy would be able to reduce oxidative stress levels by as much as 53%. This tremendous reduction supports the existing studies in the literature of the functionality of PNPLA3 1148M variants, and establishes a correlation between lipid accumulation and oxidative stress in fatty livers and alcoholic livers. The study on functionality of PNPLA3 gene through measuring the oxidative stress and damage that occurs with the liver contributed to the body of knowledge surrounding the PNPLA3 gene. The purpose of the paper is to determine the exact correlation between oxidative stress and the PNPLA3 variants. Furthermore, immortalized human hepatoma Huh-7 cell lines were used because the cell lines are frequently used as a precursor to collect data before clinical studies are approved. With further research and more data being gathered through research processes like this research project, PNPLA3 gene as a gene therapeutic method will eventually be approved for clinical trials and once it is approved to be a therapeutic options for patients with NAFLD and ALD and will help millions of Americans and people around the world suffering from this disease.

## Future Directions

This study yields future research for the advancement of PNPLA3 gene therapy. Further studies into the functionality of PNPLA3 is still needed. While ROS levels and oxidative stress have been identified and quantified across all the variants, more testing like qPCR analysis of PNPLA3 gene will be needed to verify molecular pathways and the genetic changes and the phenotypic manifestation that occurs with PNPLA3 targeted gene therapy. Another avenue of research would be testing more markers of oxidative stress like 4-HNE analysis which detects secondary byproducts of lipid induced oxidation to further analyze the oxidative stress levels between the PNPLA3 I148M variants. Finally, healthcare facilities and research institutes with access to liver tissues from patients with NAFLD and ALD can use human hepatocytes in order to measure the benefits of PNPLA3 gene therapy in actual human cells.

## Supporting information

https://docs.google.com/document/d/1H3qQhL4dBybxm_kXrR7dVIPP8ADepgYBwSsKffcIbPo/edit?usp=sharing

## Acknowledgement

I thank the Dong Lab in Indiana University School of Medicine Department of Biochemistry and Molecular Biology for providing a major contribution to this study. I thank Dr. Kim Hyeong Geug for providing the mentorship and advice crucial to this study, providing instructions for various lab techniques, and providing me with the Huh-7 cell line used in this study

