## Supplementary material for "Measurement of Oxidative Stress in Huh-7 Cell Line to Determine the Effectiveness of PNPLA3 Targeted Gene Therapy for Mitigation of Lipid and Alcohol Induced Oxidative Stress in the Liver": https://docs.google.com/document/d/1H3qQhL4dBybxm_kXrR7dVIPP8ADepgYBwSsKffcIbPo/edit?usp=sharing

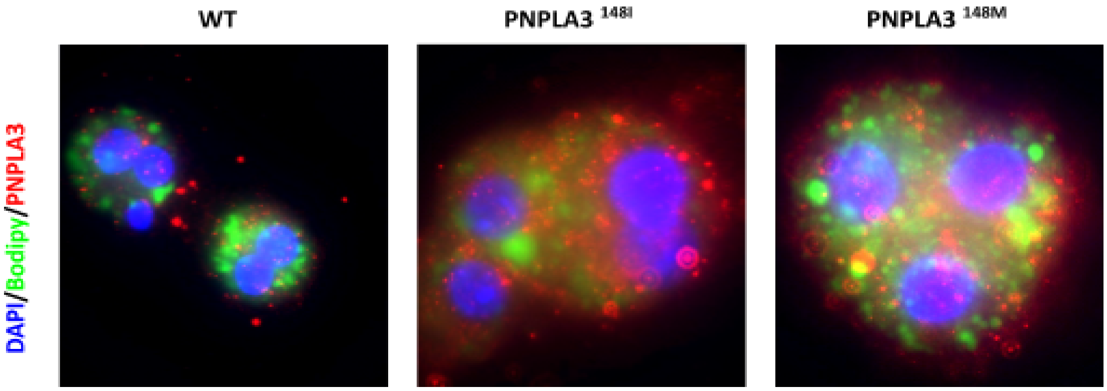


Figure 1. Genotyping fluorescence imaging data: the PNPLA3 overexpressed cell line had increased red expression and PNPLA3 148M overexpressed cell line showed increased green expression, confirming gene transfection

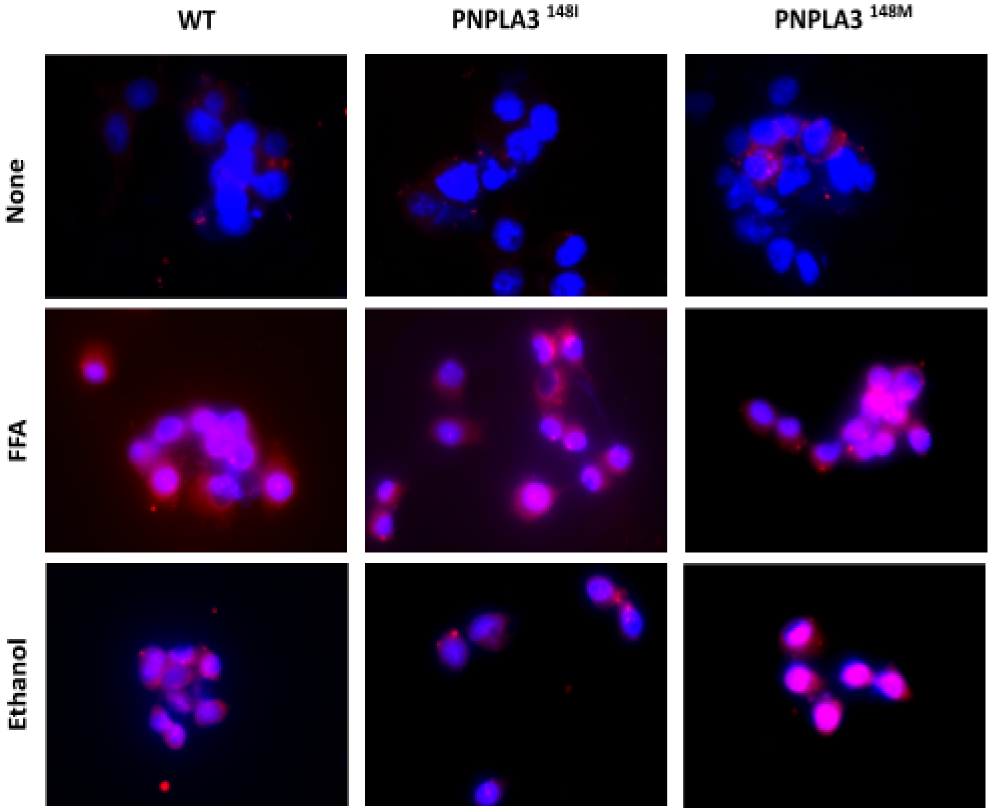


Figure 2. MitoSOX fluorescence imaging data: 148M overexpressed cell line displayed more red

fluorescence and less mitochondria compared to other groups

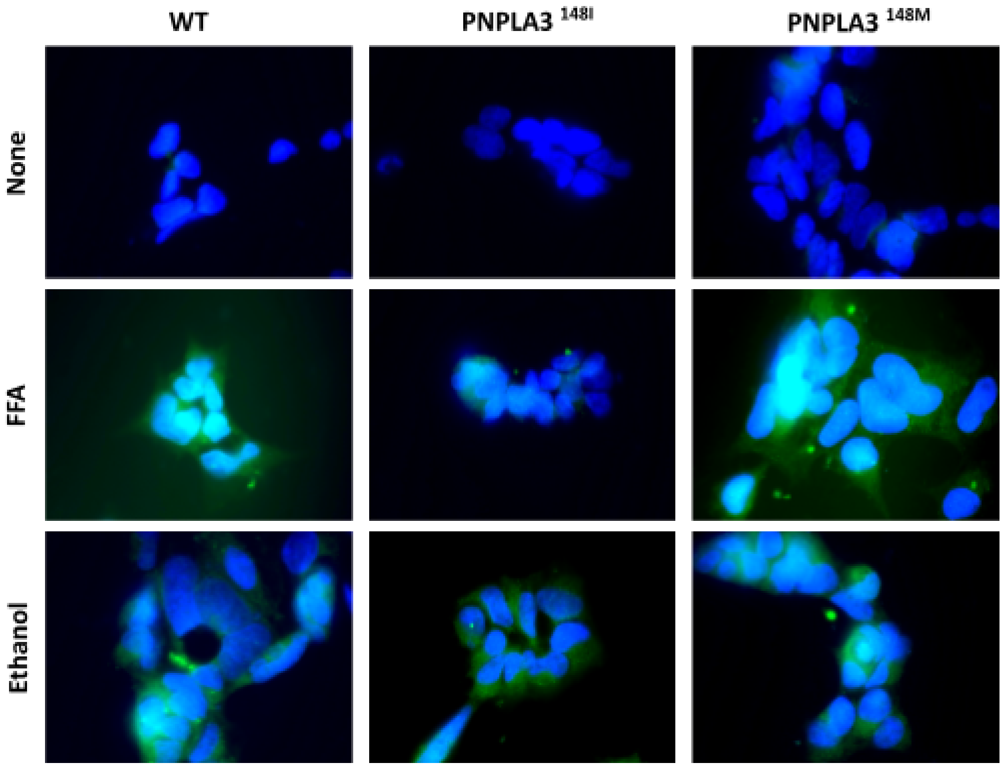


Figure 3. DCFDA fluorescence imaging data: 148M group displays the most among of greenish yellow fluorescence as compared to other groups

| Treatment / Fluorescence Intensity (MitoSOX) | WT | $PNPLA3^{148I}$ | $PNPLA3^{148M}$ |
|---|---|---|---|
| None (Two Trials; Avg) | 1.486, 1.499; 14825 | 2.120, 2.978; 2.549 | 3.226, 2.812; 3.069 |
| FFA (Two Trials; Avg) | 1.623, 1.409; 1.516 | 2.019, 1.873; 1.946 | 2.248, 2.938; 2.593 |
| Ethanol (Two Trials; Avg) | 1.672, 1.623; 1.6475 | 3.871, 5.002; 4.4365 | 3.724, 4.309; 4.0165 |

| Fluorescence Intensity (DCFDA) / Treatment | WT | $PNPLA3^{148I}$ | $PNPLA3^{148M}$ |
|---|---|---|---|
| None (Two Trials; Avg) | 1.647, 1.502; 1.5745 | 2.243, 2.697; 2.47 | 3.679, 3.226; 3.4525 |
| FFA (Two Trials; Avg) | 1.514, 1.504; 1.509 | 1.963, 2.312; 2.1175 | 2.812, 2.301; 2.5565 |
| Ethanol (Two Trials; Avg) | 1.762, 1.613; 1.6875 | 4.463, 3.877; 4.17 | 4.281, 4.412; 4.3465` |

| Fluorescence Intensity (Both DCFDA and MitoSOX) / Treatment | WT | $PNPLA3^{148I}$ | $PNPLA3^{148M}$ |
|---|---|---|---|
| None | 1.5285 | 2.5059 | 3.26075 |
| FFA | 1.5125 | 2.03175 | 2.57475 |
| Ethanol | 1.6675 | 4.30325 | 4.1815 |

Figure 4.

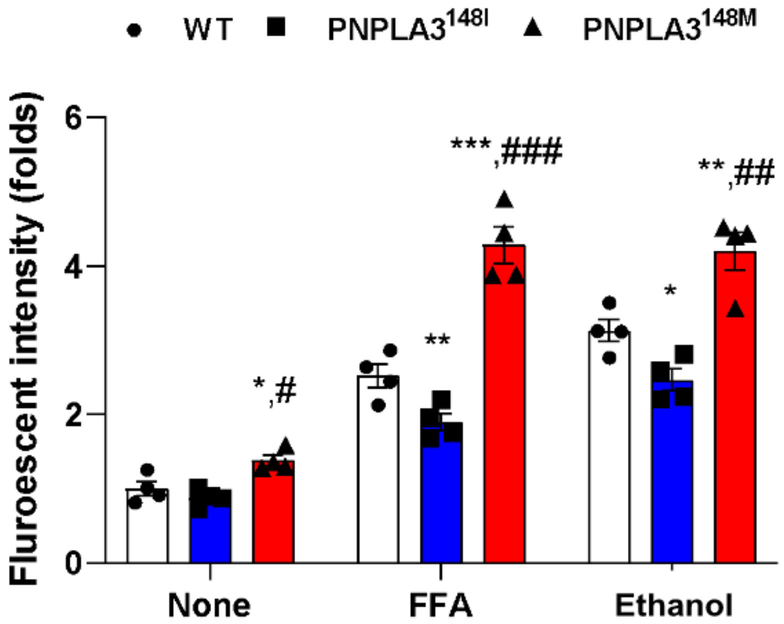


Figure 5. The fluorescence intensity as shown in this bar graphs is organized into folds, which is in a logarithmic scale, therefore, fluorescent intensity increase from 0 to 2 would be $10^2$ percent increase, and from 0 to 4 would be $10^4$ percent increase.

P values were also added, * on 148I variant bars show the p value between WT and 148I variant, * on 148M variant bars show p value between WT and 148M variant, and # shows the p value between 148I and 148 M variant. * or # represents p value less than 0.05, ** or ## represents p value less than 0.01, and *** or ### represents p value less than 0.001
